# How do Research Faculty in the Biosciences Evaluate Paper Authorship Criteria?

**DOI:** 10.1101/069468

**Authors:** Timothy Kassis

**Affiliations:** Department of Biological Engineering, Massachusetts Institute of Technology, Cambridge, Massachusetts, United States of America

## Abstract

Authorship of peer-reviewed journal articles and abstracts has become the primary currency and reward unit in academia. Such a reward is crucial for students and postdocs who are often under-compensated and thus highly value authorship as an incentive. While numerous scientific and publishing organizations have written guidelines for determining author qualifications and author order, there remains much ambiguity when it comes to how these criteria are weighed by research faculty. Here, we sought to provide some initial insight on how faculty view the relative importance of 11 criteria for scientific authorship. We distributed an online survey to 564 biomedical engineering, biology, and bioengineering faculty members at 10 research institutions across the United States. The response rate was approximately 18%, resulting in a final sample of 102 respondents. Results revealed a consensus on some criteria, such as time spent conducting experiments, but there was a lack of agreement regarding the role of funding procurement. This study provides quantitative assessments of how faculty members in the biosciences evaluate authorship criteria. We discuss the implications of these findings for researchers, especially new graduate students, to help navigate the discrepancy between official policies for authorship and the contributions that faculty truly value.

## Introduction

Authorship on peer-reviewed journal articles and conference abstracts has become the primary currency in academia and a core metric for assessing intellectual productivity and output. Thus, determining who receives authorship and ranking in the authorship list are necessary for responsible science. In particular scientific disciplines, such as mathematics, for example, authors are listed in alphabetical order. In the biosciences, however, there is a heavy emphasis on rank in the authorship list, and author order is thought to correspond to the significance of one’s contribution [1]. The last author is typically the senior author and is the principal investigator overseeing the lab, while the first author is the researcher, such as the student, postdoc or research scientist, that led the project and carried out the majority of the experimental work and manuscript preparation. Unfortunately, the authorship list has become highly politicized, where inventorship and authorship are drifting apart [2]. Determining who should be listed as the first author is usually not very difficult, but issues arise when there are multiple people involved in a study with various levels and types of contributions [3]. A recent survey showed that almost two-thirds of authors do not entirely agree with their defined contribution as indicated on journal submission disclosure forms [4]. New graduate students and researchers joining research laboratories are often unclear of the criteria that their immediate supervisors value in determining authorship. Using quantifiable metrics may help elucidate expectations for authorship on both sides.

Numerous universities, including Stanford [5], Georgia Tech [6], and Harvard University [7] have made efforts to write internal guidelines defining authorship. The majority of scientific and engineering-based organizations have also proposed guidelines describing what constitutes an author and the type of contribution required [8]. And while some attempts have been made to implement these guidelines practically within the health and biosciences [9], the criteria remain ambiguous and do not reflect which are most valued in determining authorship and rank. Biomedical journals mostly refer to guidelines set forth by the International Committee of Medical Journal Editors (ICMJE) [10]. Although these guidelines help define author inclusion, they are not of particular help when deciding authorship order. ICMJE recommends that an author meet the following four criteria:

1. Substantial contributions to the conception or design of the work; or the acquisition, analysis, or interpretation of data for the work; AND
2. Drafting the work or revising it critically for important intellectual content; AND
3. Final approval of the version to be published; AND
4. Agreement to be accountable for all aspects of the work in ensuring that questions related to the accuracy or integrity of any part of the work are investigated and resolved.

In most cases, the principal investigator of a research group is the final arbiter in determining authorship inclusion and order. In this study, we aim to break down these criteria further to elucidate how faculty in the biosciences weigh them in determining both inclusion as an author and authorship order. We do not attempt to assess these criteria from an ethical perspective [11], but rather seek to provide quantitative insight that will improve new researchers’ understanding of faculty expectations within a research project.

## Materials and Methods

There are many types of contributions in any collaborative research study. While there is no clear consensus on how to classify these contributions, we devised 11 explicit criteria based on prior literature [12] and our subjective assessment that we believe the biosciences community deems important for both determining one’s recognition as an author and their rank on the authorship list. The 11 criteria are:

1. ***Total time spent on a project***: This refers to the total amount of time devoted to the research study. Including conducting literature searches, planning experiments, performing experiments, analyzing data, writing and proofreading the manuscript.
2. ***Time spent carrying out background research and literature review***: This refers to intellectual efforts put into initially deciding on a certain research area and reviewing the literature see what has been previously accomplished in the field.
3. ***Contribution to hypothesis and idea generation***: This refers to the hypothesis upon which a study is grounded in hypothesis-driven research, or the idea for non-hypothesis-driven research such as methodologies, tools, and exploratory studies.
4. ***The contribution of a special reagent, material, or computer code***: This refers to unique material-based contributions, like a particular genetically modified cell strain, a synthesized molecule or computer code for analysis or processing.
5. ***The extent of involvement in obtaining research funding***: This refers to the process of fundraising—through writing grant proposals to funding agencies or industry collaborators.
6. ***Time spent doing experiments***: This refers to the total time conducting the experiments, whether they are simulations as part of a computational project or lab time spent culturing cells or working with animals.
7. ***The uniqueness of experimental skills and techniques:*** This refers to laboratory-based skills that are unique and require considerable prior knowledge or experience. For example, certain rodent surgical skills may take a very long time to acquire and perfect. Other skills, such as changing cell media, would not fit into this category.
8. ***Time spent analyzing data:*** This refers to taking raw data, compiling it, analyzing it, performing statistical analysis and presenting it in visual or textual formats.
9. ***Contribution to written manuscript:*** This refers to creating an outline, assembling any figures, and drafting the manuscript.
10. ***The quality of written contribution to the manuscript***: This refers to how some individuals are more efficient at writing than others, so it is difficult to assess written contributions without also evaluating the quality of one’s writing. Writing quality includes being able to explain research findings well, and employ good grammar and spelling, good structure, and flow.
11. **Time spent editing and proofreading manuscript**: This refers to the final step before submission to a journal when the lead author sends the manuscript to all the listed authors for final commenting, editing, and proofreading.

To assess the value that research faculty assign to each of these criteria, an online survey was emailed to research faculty in biology, biomedical engineering, and bioengineering at 10 research institutions (Table 1). The institutions were chosen to represent a wide geographical area of the United States with a range of research interests across the biosciences. Given the interdisciplinary nature of biological research, we decided to lump three different departments together. These departments were biology, biomedical, and bioengineering, noting that biomedical and bioengineering departments are considered the same. The survey was individually addressed and emailed to 564 total faculty members, and we received 102 responses for a total response rate of 18.1%. Faculty members were identified by their listings on their department’s website. Eligible faculty were those who provided an email address on their university departmental page. No other criterion for sample selection was used. No follow-up reminders were sent, nor were other optimization strategies used for the purpose of this study [13]. The survey was designed to be straightforward and easy to fill out to maximize the response rate. For each criterion, the respondent had to indicate, on a scale of 1-10, how important they thought the criterion was, with one being the least important and ten being the most important. Specifically, we asked:

> *“On a scale of 1-10, how important are the following factors in determining authorship and authorship rank on a peer-reviewed journal paper? (Please note this only applies to life sciences/biosciences/biomedical engineering).”*

Only the main title of each criterion listed above in bold was provided to the survey respondents. The descriptions listed under each criterion are strictly for clarification purposes for the readers of this manuscript. Survey results are provided as a .csv supplemental file (**S1 File**).

**Table 1:** Institutions and departments the survey was sent to.

| **Institution** | **Department/Program** | **# of Faculty Contacted** |
| --- | --- | --- |
| University of California - Los Angeles | Bioengineering | 29 |
| Georgia Institute of Technology | Bioengineering | 98 |
| Johns Hopkins University | Biomedical Engineering | 62 |
| Duke University | Biomedical Engineering | 72 |
| University of California - Davis | Biomedical Engineering | 24 |
| University of Texas - Austin | Biomedical Engineering | 32 |
| Texas A&M University | Biomedical Engineering | 25 |
| University of Washington | Biology | 69 |
| Stanford University | Biology | 55 |
| Arizona State University | Biology | 98 |
| **Total:** |  | **564** |
| **Responses (Response rate):** |  | **102 (18.1%)** |

## Statistical Analysis

A D’Agostino & Pearson omnibus normality test was conducted, and only three of the 11 criteria were normally distributed. Thus in our results, we focus on the median, the 25^th^ and 75^th^ quartiles for data reporting as they are more appropriate for skewed, non-normal distributions. We also use the coefficient of variation as a metric to explain the dispersion in the response histograms. All graphing and analysis was carried out using Graphpad Prism 6 (GraphPad Software Inc. La Jolla, CA, USA).

## Results and Discussion

Many criteria are used to assess authorship. We found the time spent performing experiments had the highest overall median importance score as assessed by our faculty respondents. The intellectual contribution of the hypothesis (for hypothesis-driven research) or coming up with a study idea had the second highest score. The very act of contributing a reagent, material or computer code— even when it is unique to the person contributing it—tied for least important with the extent of one’s involvement in obtaining funding. Overall, there seemed to be a consensus that the time spent conducting experiments, coming up with a hypothesis, analyzing data, and writing the manuscript were the four most important criteria for both determining one’s authorship status and rank (Fig 1A). The total time spent on a project was assessed as being important, but 19.6% of the respondents selected a neutral score of 5 indicating that, by itself, time spent on a project should not necessarily factor into authorship, which might reflect the fact that time alone does not translate to productivity. The median value was 7/10, and this criterion ranked fifth of all contributions (Fig 1B).

**Fig 1.**
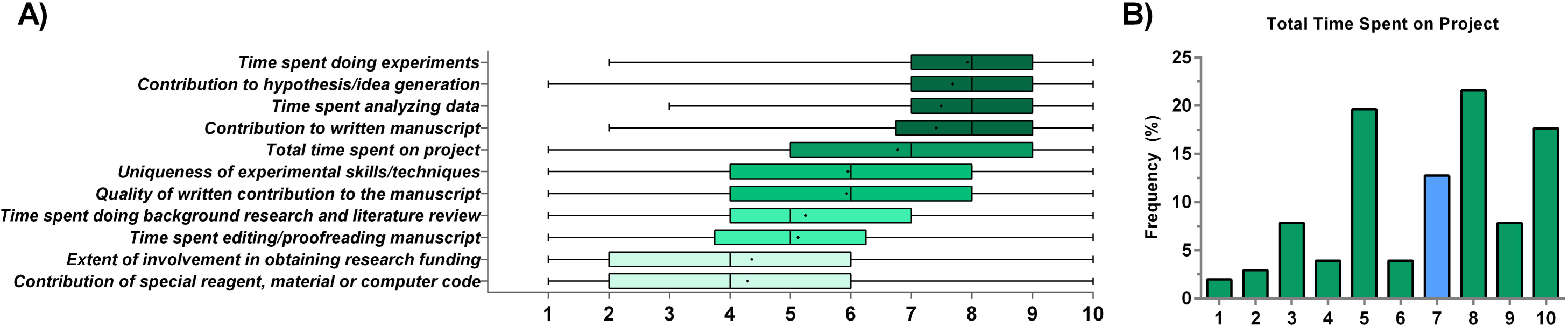
A ranking of the relative importance of 11 authorship criteria. A) Ranking of various criteria according to research faculty in the biomedical sciences. The middle line represents the median, the edges of the box represent the 25% and 75% quartiles; the whiskers represent the range and the ‘+’ mark is the mean. B) Total time spent has a high weight, but the majority of faculty do not appear to hold a strong opinion about this as reflected by the 19.6% of respondents giving it a score of 5, probably due to the fact that time spent does not necessary equate with an intellectual contribution to the study. N = 102 for all data presented. The blue bar represents the scale median.

The scores did not follow a normal distribution, so we calculated the coefficient of variation (CV) as the main metric for quantifying spread instead of the standard deviation (Fig 2). While all criteria had a CV greater than 23%, two criteria stood out as having a very high CV. The first, with the highest CV of 57%, was the extent of involvement in obtaining funding, while the second highest CV with 53% was the contribution of a material. The four criteria with the highest median scores (Fig 1A) also had the lowest CV values.

**Fig 2.**
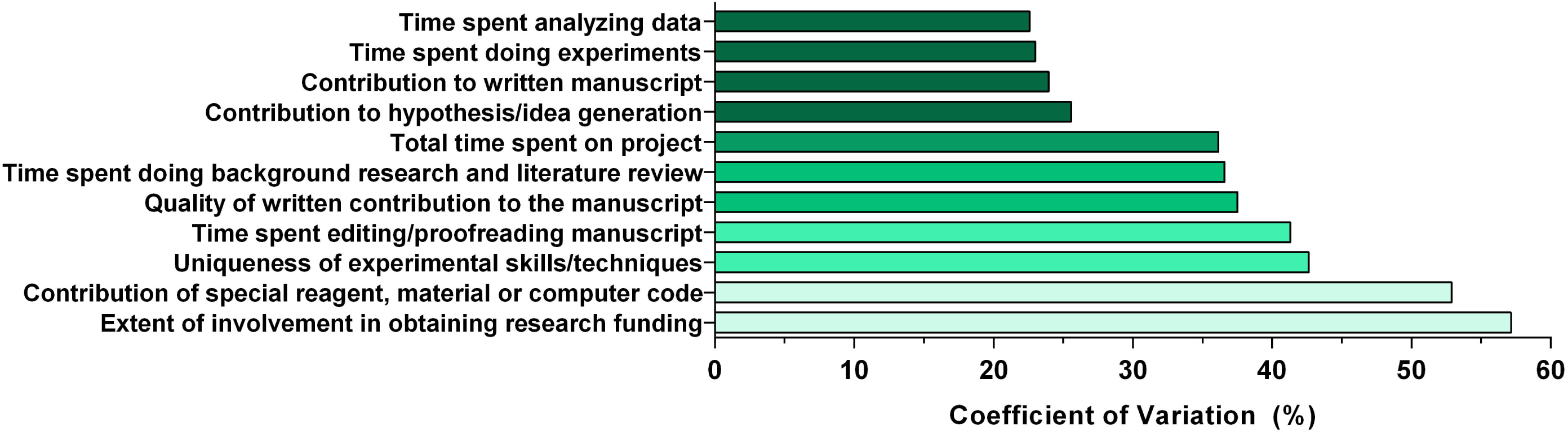
Quantifying spread as a metric for agreement. Coefficient of variation (CV) used as a metric to quantify spread of the scores. Lower values represent a higher agreement between faculty on the importance score. Higher values imply the distribution was highly dispersed and there was little consensus regarding the score.

Figure 3 presents histograms of the responses describing four criteria that revolve around preparing for the study. These include background research, hypothesis generation, the contribution of various material goods, and obtaining funding. We found that almost 23% of respondents had a neutral opinion of the value of background research, such that this alone did constitute direct involvement with the study. The median was 5/10 (Fig 3A). There was a clear consensus regarding the importance of generating a hypothesis or idea with the majority of faculty who weighted this criterion strongly with a median of 8/10 and the majority giving it a 10/10 score (Fig 3B). The contribution of a special reagent, material, or computer code did not constitute grounds for authorship, with almost 23% of faculty giving it a 2/10 score with a median of 4/10 (Fig 3C). An issue of great controversy in academia is whether involvement in obtaining funding, such as writing a grant proposal, justifies authorship. Although the median was 4, there was no clear consensus on the importance of this criterion as indicated by both the low median (Fig 3D) and high CV (Fig 2).

**Fig 3.**
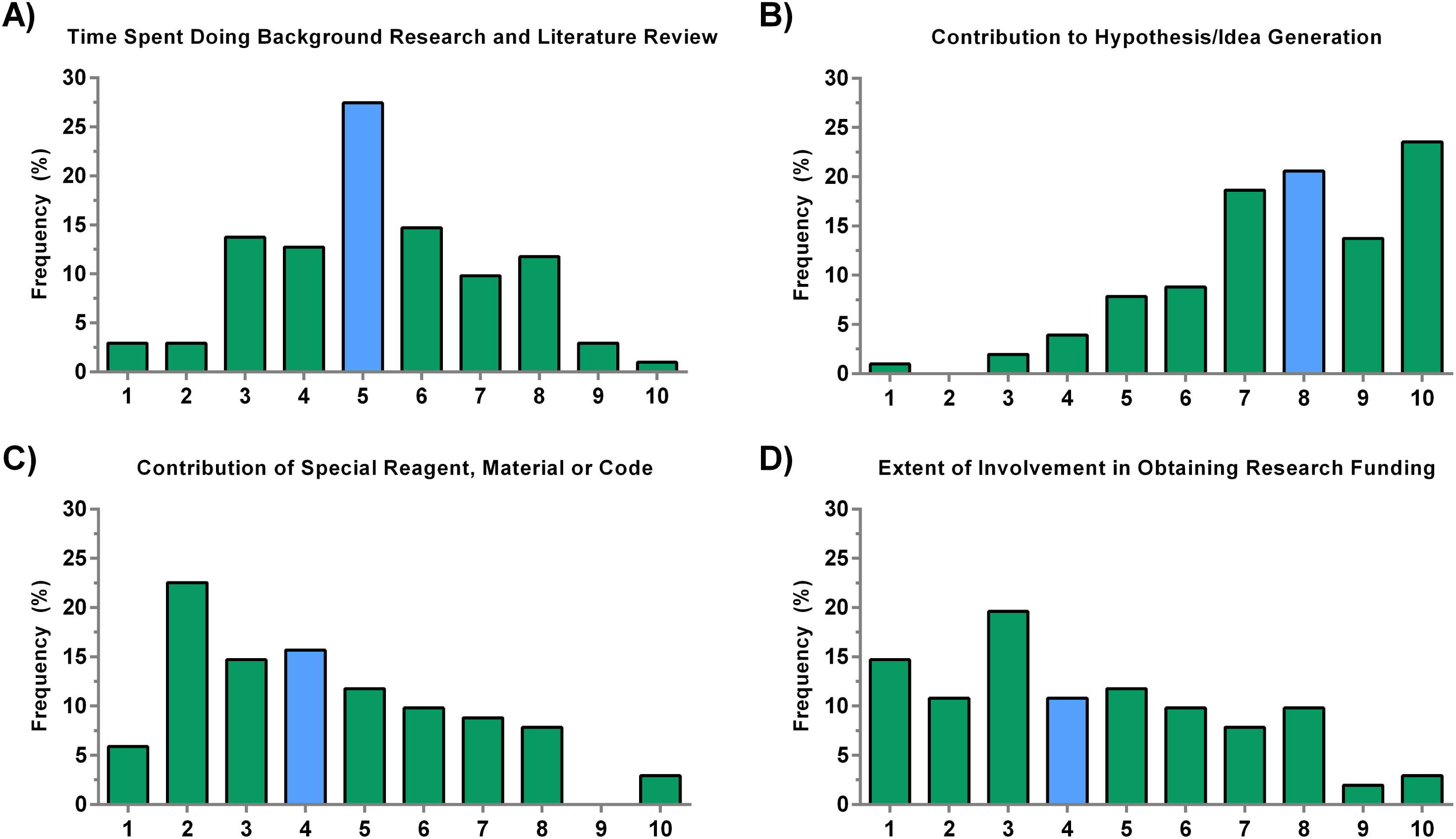
**Criteria involved in preparation for a research study.** A) A majority of responses regarded time spent doing background research as neutral. B) The histogram is clearly skewed, indicating that contribution to the hypothesis and initial idea is crucial. C) Although it is difficult to generalize to all material-based contributions, our survey respondents lean towards the idea that contributing a special reagent, material, or computer code alone does not justify authorship eligibility and rank. D) There is no clear consensus on the role that obtaining funding plays. N=102. The blue bars represent the median.

The next two criteria involved experimental aspects of the project. These included time spent performing experiments as well as any required skill or technique. We found that respondents highly value the time put into performing experiments with a median score of 8/10, and almost 22% of faculty scored it as 10/10 (Fig 4A). Some experiments require special skills that involve more than simply following a protocol. These can include surgical techniques, specific cell handling procedures, and skills that would typically require a lot of experience to adequately master or which cannot easily be replaced. Possibly due to the vague nature of these skills and diverse techniques employed across the biosciences, there was no clear consensus on the importance of this criterion. The median was 6/10 (Fig 4B), but the scores had a high CV of 43%.

**Fig 4.**
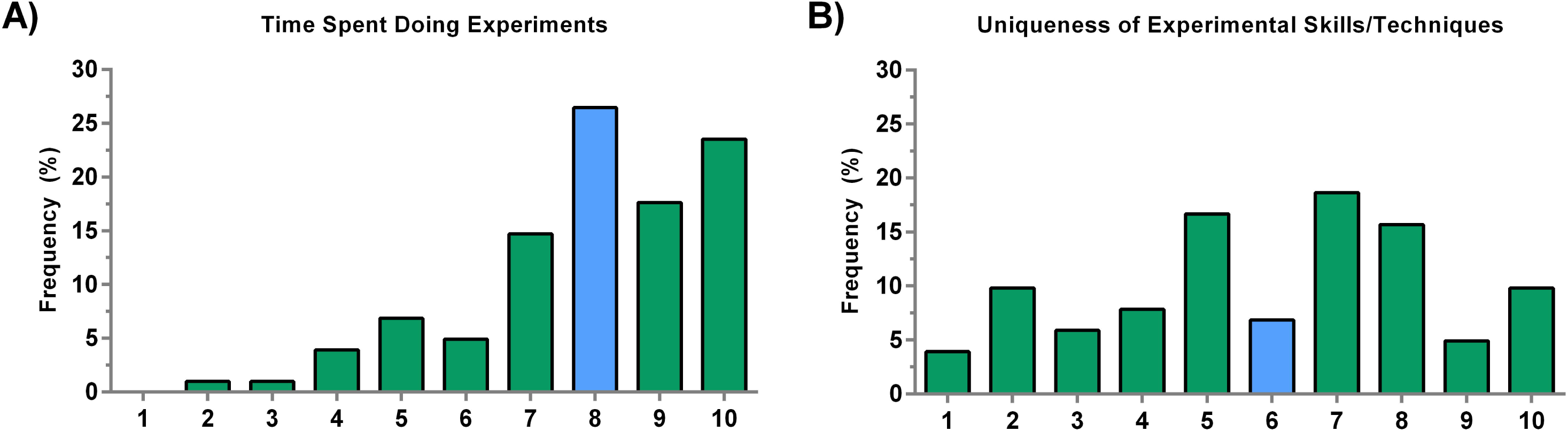
Criteria applicable during the experimental aspects of a research study. A) Time spent conducting experiments is important with the majority of faculty scoring it high. B) The uniqueness of experimental skills and techniques had a median of 6 with a high coefficient of variation. The blue bars represent the median.

Last but not least, we evaluated the four criteria that corresponded with the post-experimental stage of a research study. These included data analysis, writing the manuscript, actual quality of the manuscript content contribution, and editing/proofreading of the manuscript. Both analyzing data and writing the manuscript were considered very important by the respondents, both of which had a median of 8/10 (Fig 5A and **5B**). Almost 25% of respondents agreed that the quality of the written content is important and gave it an 8/10 score, but there was significant variation in responses, resulting in a median of 6/10 (Fig 5C). The final step of the paper submission, which involves final edits and proofreading, had a median of 5/10, again indicating that most respondents do not necessarily have a strong opinion about this criterion (Fig 5D).

**Fig 5.**
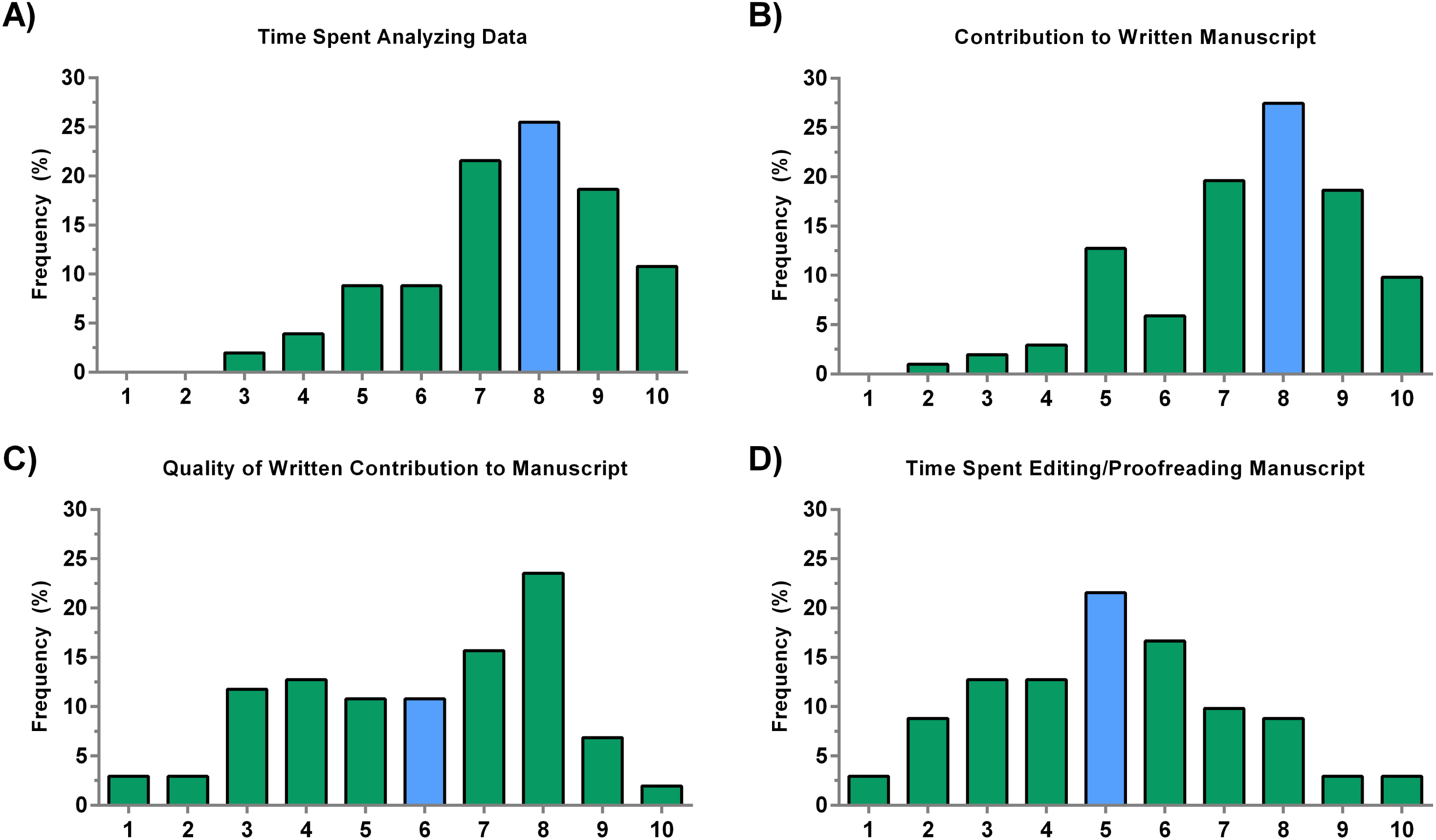
Criteria representing post-experimental stage of a research study. A) The amount of time spent analyzing data is as important as writing the manuscript (B). C) The quality of the contribution to the written manuscript is also important, but there were several respondents that indicated that it was not. D) The majority of faculty had a neutral stance regarding the time spent editing and proofreading the manuscript. The blue bars represent the median.

As scientific studies become more interdisciplinary and collaborative in nature, authorship lists have grown. Here, we attempted to quantify how faculty in the biosciences value various academic contributions. We hope that these results remove some of the ambiguity surrounding this important issue and provide some insight for new graduate students and researchers. We believe that the data presented here will serve three main goals; 1) provide authorship decision makers, such as the first and senior author, explicit criteria that will assist them when crediting contributions on their manuscripts. 2) give researchers, especially new graduate students, metrics that they can use when discussing their role in the project, and 3) better understand some aspects of authorship attribution in the biosciences academic setting. While our study provides some quantifiable evidence on the attitudes held by faculty at research institutions, it is bounded by several limitations. Some research projects may not involve all 11 criteria, so if a certain criterion was ranked low, it could be that it just does not exist within certain types of research, and hence faculty respondents working in that research area might have scored it low or given it a neutral score. This may account for some of the high CV values apparent in the results. It is also important to mention that due to the small sample size, the scores here may not be representative of all faculty in the biosciences community, and a much more comprehensive study is necessary to arrive at generalizable conclusions. Additionally, since no survey reminders or follow-up was conducted after the initial email, it is likely that there was a higher likelihood of selection bias for faculty with a greater tendency to have certain opinions. The survey was also sent to all email accessible faculty members in the same departments, so there was the potential for them to discuss questions before returning the survey. While we believe the findings presented here illuminate good initial results, they merely provide a basis for much more comprehensive and systematic surveys that should be conducted.

## Conclusions

While the data gained through this survey is limited in scope, such information helps advance a standardized method for assessing authorship inclusion and rank on the authorship list and begins to understand how members of the biosciences community faculty evaluate various criteria. We hope that in the future, objective methodology can standardize authorship across research laboratories and identify where author contributions can be better defined and tracked. In this small study, we provided some initial quantifiable insight to help early researchers and the biosciences community as a whole, but more work by individual researchers, organizations, and publishers is needed to arrive at generalizable and clearly communicated criteria for determining publication authorship and rank.

## Acknowledgements

I would like to thank all the faculty respondents for answering the survey. I also thank all faculty members who had direct input to give especially Dr. Henry Sauermann of the Georgia Institute of Technology for providing input on an early version of this manuscript and providing constructive criticism. During my Ph.D. research, I was fortunate enough to be working in a lab where authorship conflicts were rarely an issue.

## Supporting Information

**S1 File. Survey Results.** Dataset generated from survey results in .csv format.

